# Production of human milk fat substitute by engineered strains of *Yarrowia lipolytica*

**DOI:** 10.1101/2021.07.30.452670

**Authors:** Govindprasad Bhutada, Guillaume Menard, Rodrigo Ledesma-Amaro, Peter J Eastmond

## Abstract

Human milk fat has a distinctive stereoisomeric structure where palmitic acid is esterified to the middle (sn-2) position on the glycerol backbone of the triacylglycerol and unsaturated fatty acids to the outer (sn-1/3) positions. This configuration allows for more efficient nutrient absorption in the infant gut. However, the fat used in most infant formulas originates from plants, which tend only to esterify palmitic acid to the sn-1/3 positions. Oleaginous yeasts provide an alternative source of lipids for human nutrition. However, these yeasts also exclude palmitic acid from the sn-2 position of their triacylglycerol. Here we show that *Yarrowia lipolytica* can be engineered to produce triacylglycerol with more than 60% of the palmitic acid in the sn-2 position, by expression of a lysophosphatidic acid phosphatase with palmitoyl-Coenzyme A specificity, such as LPAAT2 from *Chlamydomonas reinhardtii*. The engineered *Y. lipolytica* strains can be cultured on glycerol, glucose, palm oil or a mixture of substrates, under nitrogen limited condition, to produce triacylglycerol with a fatty acid composition that resembles human milk fat, in terms of the major molecular species; palmitic, oleic and linoleic acids. Culture on palm oil or a mixture of glucose and palm oil produced the highest lipid titre in shake flask culture and a triacylglycerol composition that is most similar with human milk fat. Our data show that an oleaginous yeast can be engineered to produce a human milk fat substitute (β-palmitate), that could potentially be used as an ingredient in infant formulas.

## 1. Introduction

Human milk is the best source of nutrition for infants and it is their main food during the first four to six months of life (Innis 2011; Wei et al., 2019). The lipid fraction provides around half the infant’s calories and consists of approximately 98% triacylglycerol (TAG) (Wei et al., 2019). In human milk fat, palmitic acid (16:0) is esterified to the middle (sn-2) position on the glycerol backbone and oleic acid (18:1) and linoleic acid (18:2) to the outer (sn-1/3) positions (Breckenridge et al., 1969; Christie and Clapperton, 1982; Giuffrida et al., 2018), giving the TAG molecules distinctive stereochemistry that clinical trials have suggested can assist nutrient absorption in the infant gut (Innis 2011; Béghin et al., 2018). However, the TAG used in most infant formulas is derived from plants (Wei et al., 2019), which exclude 16:0 from the sn-2 position (Brockerhoff and Yurkowsk, 1966; Christie et al., 1991). Mixtures of vegetable fats (plus algal and fish oils) can be blended to mimic the fatty acid (FA) composition of human milk, but not its stereoisomeric structure (Wei et al., 2019).

To address this issue, companies have developed a class of ‘structured lipids’ called human milk fat substitutes (HMFS) (or β-palmitate) that are produced by enzyme-catalyzed acidolysis (or alcoholysis and esterification) of fractionated vegetable TAG and FAs using sn-1/3 regioselective lipases (Wei et al., 2019). These HMFS (e.g. Betapol and InFat) currently provide enrichment of 16:0 at the sn-2 position of up to 60% in the final fat phase of infant formulas. HMFS are relatively costly compared to conventional vegetable fats and it remains technically challenging to manufacture a true mimetic at industrial scale that is affordable (Ferreira-Dias and Tecelão, 2014). HMFS ideally requires more than 60% of the 16:0 to be esterified to the sn-2 position in the final fat phase to replicate human milk fat (Breckenridge et al., 1969; Lopez-Lopez et al., 2002). Indeed, the main TAG species in human milk are 1,3-dioleoyl-2-palmitoyl-glycerol (OPO) and 1-oleoyl-2-palmitoyl-3-linoleoyl-glycerol (OPL) or LPO (Giuffrida et al., 2018) and formula makers would likely benefit from an affordable source of pure OPO and OPL that they could then blend with cheaper vegetable fats. A substantial enrichment of 16:0 at the sn-2 position is also found in other animal fats used for human nutrition such as butterfat (Christie and Clapperton, 1982) and lard (Christie and Moore, 1970). Hence, HMFS-type structured lipids also have potential markets in meat and dairy substitute products.

We are investigating whether oleaginous organisms, that normally exclude 16:0 from the sn-2 position of their TAG, can be engineered to produce HMFS. Using the model oilseed plant *Arabidopsis thaliana*, we have recently showed that it is possible to produce TAG with more than 80% of the 16:0 esterified to the sn-2 position and where around 40% of the molecules are OPO (van Erp et al., 2019; 2021). TAG is formed by a cytosolic glycerolipid biosynthetic pathway that’s situated on the endoplasmic reticulum (ER) in plants and the enzyme responsible for acylation of the sn-2 position is lysophosphatidic acid acyltransferase (LPAT) (Ohlrogge and Browse, 1995). ER-localized LPAT isoforms discriminate against a 16:0-Coenzyme A (CoA) substrate in plants (Kim et al., 2005). By expressing LPAT isoforms with a preference for 16:0-CoA from other organisms, we were able to drive incorporation of 16:0 into the sn-2 position of TAG, both in *A. thaliana* with a wild type FA composition (van Erp et al., 2019), and in a multi-mutant with high 16:0 and 18:1 content in its seeds (van Erp et al., 2021).

However, to achieve a sufficient enrichment of 16:0 at sn-2 (above 60%) in *A. thaliana* seed oil, we found that suppression of endogenous *LPAT2* expression (Kim et al., 2005) and disruption of phosphatidylcholine:diacylglycerol cholinephosphotransferase (PDCT) (Lu et al., 2009) were also required. PDCT is a plant-specific enzyme that catalyzes head group exchange between diacylglycerol (DAG) and phosphatidylcholine (PC) in *A. thaliana* seeds (Lu et al., 2009; Bates et al., 2012). PC is the site of acyl-lipid desaturation (Miquel and Browse, 1992) and acyl editing in plants (Stymne and Stobart, 1984; Wang et al., 2012; Bates et al., 2012) and so disruption of PDCT may limit loss of 16:0 from the sn-2 position prior to TAG synthesis (van Erp et al., 2019).

In this study we decided to investigate whether HMFS can be produced by an oleaginous yeast. We chose to use *Yarrowia lipolytica* as a host. *Y. lipolytica* grows on a variety of carbon sources (Groenewald et al., 2014; Spagnuolo et al., 2018), it is amenable to genetic engineering (GE) and it has a history of use by industry (Groenewald et al., 2014; Ledesma-Amaro and Nicaud, 2016). *Y. lipolytica* biomass has been defined as a safe novel food by the European Food Standards Agency (Turck et al., 2019) and the US Food and Drug Administration has granted Generally Recognized As Safe status to several products made ‘with the assistance of’ GE *Y. lipolytica* strains (Groenewald et al., 2014), including specialty lipids intended for use in human nutrition such as TAG containing omega-3 long chain polyunsaturated FAs (Xue et al., 2013; GRN 000355). Substantial efforts have also been made to improve *Y. lipolytica* lipid productivity using GE (Ledesma-Amaro and Nicaud, 2016) and strains have been reported that achieve cell lipid content, titer, productivity and yield of up to ∼70%, 99g L^-1^, ∼1.2 g^-1^ L^-1^ h^-1^ and ∼0.27 g g^-1^ in nitrogen-limited glucose-fed batch culture (Qiao et al., 2017). The FA composition of *Y. lipolytica* strains cultured on glucose (Carsanba et al., 2020) already resembles that of human milk fat, in terms of the major FA molecular species 16:0, 18:1 and 18:2 (Breckenridge et al., 1969; Giuffrida et al., 2018). However, yeasts (including *Y. lipolytica*) are known to exclude 16:0 from the sn-2 position of their TAG (Thorpe and Ratledge 1972) and so reversing this selectivity is a key step to producing a HMFS in these oleaginous organisms.

## 2. Materials and methods

### 2.1. Material and growth conditions

The *Yarrowia lipolytica* strains used in this study are listed in Table 1. The media and growth conditions for *Escherichia coli* and *Y. lipolytica* have been described by Sambrook and Russell (2001) and Barth and Gaillardin (1996), respectively. *Y. lipolytica* cultures were grown in nitrogen limited minimal media as described previously (Bhutada et al., 2017), except that palm oil emulsified with 1% (v/v) Tween 80 was used as a carbon source in some experiments. For the growth of *ura3Δ* or *leu2Δ* auxotrophic strains, media were supplemented with 0.1 g L^-1^ uracil or leucine. All PCR reactions for cloning and amplification of sequencing templates were performed using Herculase II Fusion DNA Polymerase (Agilent Technologies), and using GoTaq (Promega) for confirmation of chromosomal integration of the transformation cassettes. The restriction enzymes used in this study were obtained from Roche or New England Biolabs (NEB). The DNA fragments from PCR and restriction digestion were recovered from agarose gels using GeneJET kits (Thermo Scientific). For ligations, the Fast-Link DNA Ligation Kit (Epicenter) or Gibson assembly (Gibson et al., 2009) was used. For transformation into *Y. lipolytica* standard protocols for lithium acetate were used (Le Dall et al., 1994). All plasmids and PCR primers are listed in Supplementary Table 1.

**Table 1.**
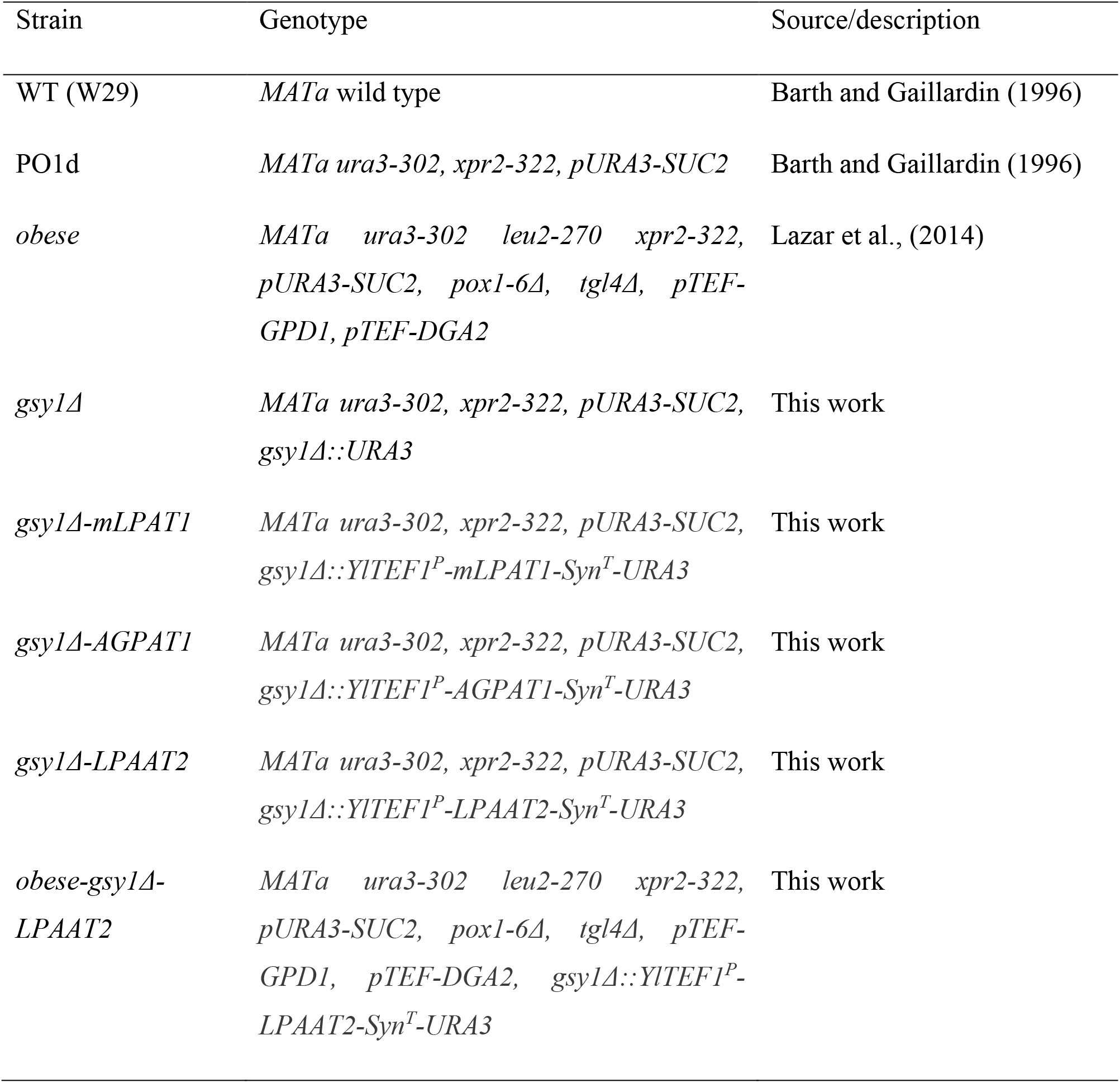
*Y. lipolytica* strains used in this study.

| Strain | Genotype | Source/description |
| --- | --- | --- |
| WT (W29) | *MATa* wild type | Barth and Gaillardin (1996) |
| PO1d | *MATa ura3-302, xpr2-322, pURA3-SUC2* | Barth and Gaillardin (1996) |
| *obese* | *MATa ura3-302 leu2-270 xpr2-322, pURA3-SUC2, pox1-6Δ, tgl4Δ, pTEF-GPD1, pTEF-DGA2* | Lazar et al., (2014) |
| *gsylΔ* | *MATa ura3-302, xpr2-322, pURA3-SUC2, gsylΔ::URA3* | This work |
| *gsylΔ-mLPAT1* | *MATa ura3-302, xpr2-322, pURA3-SUC2, gsylΔ::YITEF1<sup>P</sup>-mLPAT1-Syn<sup>T</sup>-URA3* | This work |
| *gsylΔ-AGPAT1* | *MATa ura3-302, xpr2-322, pURA3-SUC2, gsylΔ::YITEF1<sup>P</sup>-AGPAT1-Syn<sup>T</sup>-URA3* | This work |
| *gsylΔ-LPAAT2* | *MATa ura3-302, xpr2-322, pURA3-SUC2, gsylΔ::YITEF1<sup>P</sup>-LPAAT2-Syn<sup>T</sup>-URA3* | This work |
| *obese-gsylΔ-LPAAT2* | *MATa ura3-302 leu2-270 xpr2-322, pURA3-SUC2, pox1-6Δ, tgl4Δ, pTEF-GPD1, pTEF-DGA2, gsylΔ::YITEF1<sup>P</sup>-LPAAT2-Syn<sup>T</sup>-URA3* | This work |

### 2.2. Cloning and transformation

To obtain *Y. lipolytica* strains expressing eukaryotic LPATs with a substrate preference of 16:0-CoA, we synthesised codon optimized versions of *Brassica napus mLPAT1* (van Erp et al., 2019; 2021), *Homo sapiens AGPAT1* (van Erp et al., 2021) and *Chlamydomonas reinhardtii LPAAT2* (Kim et al., 2018) together with the Tsynt25 (Syn^T^) synthetic terminator fragment (Curran et al., 2015) (Supplementary Figure 1). We then linearised the plasmid pGMKGSY_12 (Bhutada et al., 2017), which harbours a glycogen storage elimination cassette flanked by ∼1 kb recombination regions for the glycogen synthase (*GSY1*) locus, with *Hind*III. We PCR amplified the strong constitutive *TEF1* promoter from *Y. lipolytica* W29 genomic DNA using the primers pair TEF-GSY-F and TEF-GSY-R and assembled the fragments by Gibson assembly to produce the plasmid pGSYTEF. The codon optimised LPAT-Syn^T^ fragments were excised from pUC7 using *Hind*III. pGSYTEF was digested with the same enzyme to linearize the vector and it was re-ligated with the gel-purified LPAT-SynT fragments to produce *pTEF-mLPAT1, pTEF-AGPAT1* and *pTEF-LPAAT2*. The correct assembly of the episomal *YlGSY1*^*P*^*-loxP-URA3-loxP-*TEF^*P*^mLPAT1Syn^T^*-YlGSY1*^*T*^, *YlGSY1*^*P*^*-loxP-URA3-loxP-*TEF^*P*^AGPAT1Syn^T^*-YlGSY1*^*T*^ and *YlGSY1*^*P*^*-loxP-URA3-loxP-*TEF^*P*^LPAAT2Syn^T^*-YlGSY1*^*T*^ cassette was confirmed by sequencing. These cassettes were excised with *Not*I, gel-purified and used for transformation of strain PO1d or *obese* (Table 1). Transformants with integration of the cassette at the *GSY1* locus were identified by Lugol’s iodine staining (1% KI, 0.5% I2) and confirmed by PCR amplification of genomic DNA flanking the integration site using primer GSY1^P^-F with mLPAT1-R, AGPA1-R, LPAAT2-R or GSY1^T^-R and by sequencing the products.

**Figure 1.**
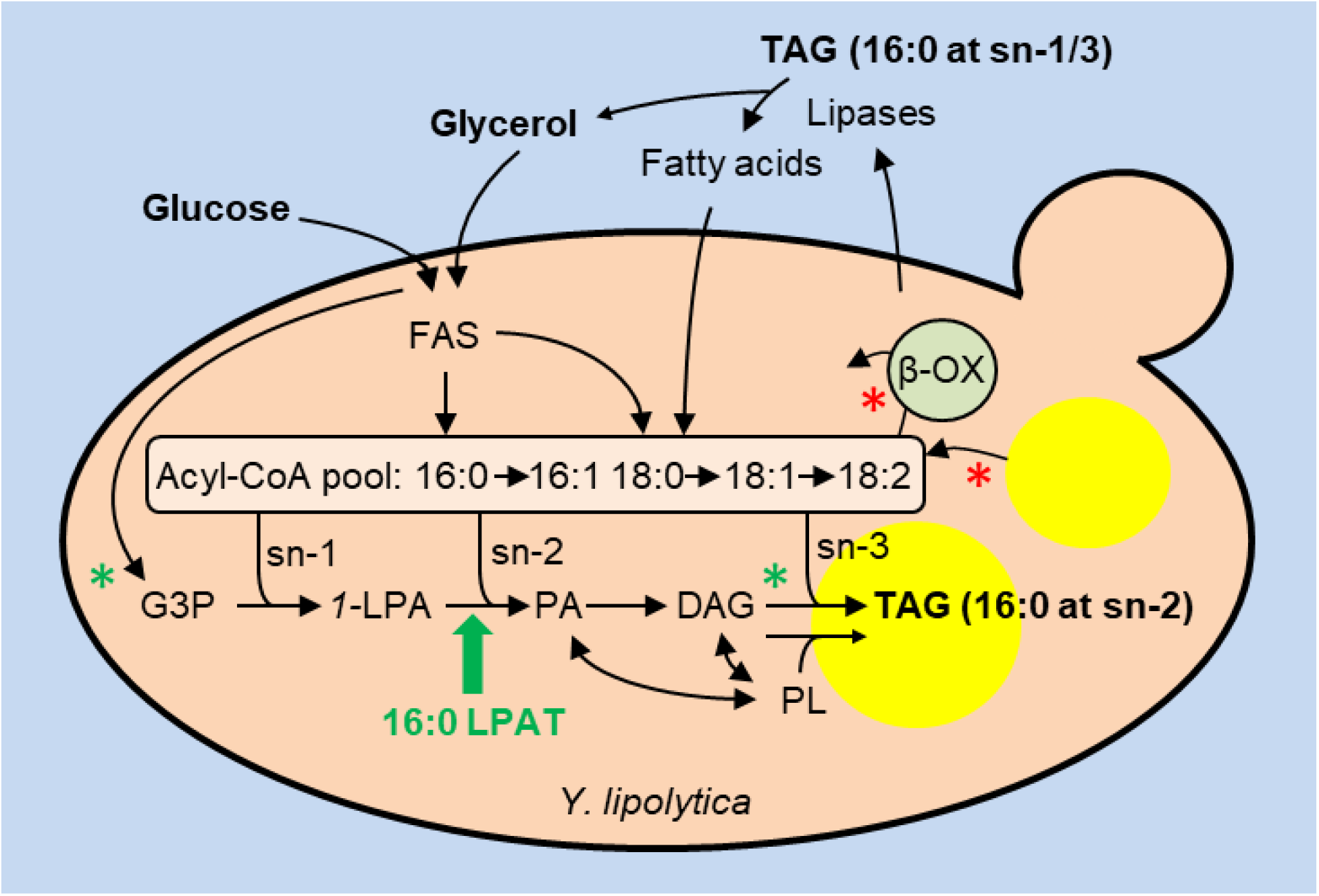
A diagram illustrating the strategy used in this study to produce HMFS in *Y. lipolytica*, using glycerol, glucose and/or palm oil as substrates. Heterologous expression of an LPAT with 16:0-CoA specificity (green arrow) was used to enable 16:0 to be esterified to the sn-2 position of *1*-LPA and the product PA is then metabolised to TAG. Green and red asterisks mark modifications in an *obese* strain (Lazar et al., 2014) used to increase lipid accumulation. FAS, fatty acid synthase; β-OX, peroxisomal FA β-oxidation; 16:0, palmitic acid; 16:1, palmitoleic acid; 18:0 stearic acid, 18:1, oleic acid; 18:2, linoleic acid; CoA, Coenzyme A; G3P, glycerol-3-phosphate; *1-*LPA, sn-1 lysophosphatidic acid; PA, phosphatidic acid, DAG, diacylglycerol, TAG, triacylglycerol; PL, major phospholipids.

### 2.3. Lipid analysis

Lipids were extracted from *Y. lipolytica* cell pellets as described by (Bhutada et al., 2017), following the methods of Hanscho et al., (2012). TAG was purified from the lipids and regiochemical analysis was performed by lipase digestion following the methods described by (van Erp et al., 2019). The total lipids, TAG and 2-monoacylglycerol fractions were trans-methylated and their FA content were quantified by gas chromatography (GC) coupled to flame ionization detection, as described previously (van Erp et al., 2019), using a 7890A GC system fitted with DB-23 columns (30 m x 0.25 mm i.d. x 0.25 µm) (Agilent Technologies). Tripentadecanoin was added to the lipid fractions prior to transmethylation to provide an internal standard for quantification.

### 2.4. Dry biomass determination

To determine dry biomass, the cell pellets from culture samples were washed twice with distilled water, filtered on 0.45 μm nitrocellulose filters, dried at 97°C overnight and weighed (Bhutada et al., 2017).

### 2.5. Statistical analyses

All experiments were carried out using three or more biological replicates and the data are presented as the mean values ± standard deviation of the mean (SD). For statistical analysis we either used one-way analysis of variance (ANOVA) with post-hoc Tukey HSD (Honestly Significant Difference) tests, or two-tailed Student’s t-tests.

## 3. Results

### 3.1. LPAT expression in Y. lipolytica produces TAG with over 60% of 16:0 at the sn-2 position

In yeasts, TAG is formed by a glycerolipid biosynthetic pathway whose enzymes are situated on the ER and on lipid droplets (LD) (Klug and Daum, 2014; Bredeweg et al., 2017) and LPATs are responsible for acylation of the sn-2 position (Benghezal et al., 2007; Jain et al., 2007; Ayciriex et al., 2012; Fig. 1). To determine whether heterologous expression of LPATs with specificity for 16:0-CoA can drive incorporation of 16:0 into the sn-2 position of TAG in *Y. lipolytica*, we selected three different LPATs of eukaryotic origin to test; LPAT1 from *B. napus* (Bourgis et al., 1999), AGPAT1 from *H. sapiens* (Agarwal et al., 2011) and LPAAT2 from *C. reinhardtii* (Kim et al., 2018). All three LPATs have been reported to be capable of using 16:0-CoA as a substrate (Bourgis et al., 1999; Agarwal et al., 2011; Kim et al., 2018). AGPAT1 and LPAAT2 are localised to the ER (Agarwal et al., 2011; Kim et al., 2018), whereas LPAT1 is situated in the chloroplast inner membrane (Bourgis et al., 1999). However, we have shown that expression of a modified version of LPAT1 (mLPAT1) that lacks the N-terminal chloroplast transit sequence results in ER localisation in plants (van Erp et al., 2019).

We synthesised *mLPAT1, AGPAT1* and *LPAAT2* cDNAs that are codon optimised for expression in *Y. lipolytica*. We then cloned them downstream of the native translational elongation factor EF-1 alpha (*TEF1*) promoter and integrated the cassettes at the glycogen synthase (*GSY1*) locus (Bhutada et al., 2017) in strain PO1d. Up to 40 Ura + colonies were selected, screened for glycogen deficiency and cassette integration at the *GSY1* locus was confirmed by PCR on genomic DNA and sequencing. Representative *gsy1Δ-mLPAT1, gsy1Δ-AGPAT1* and *gsy1Δ-LPAAT2* strains, together with wildtype strain W29 (WT) and *gsy1Δ* controls, were grown in triplicate in shake flask cultures in nitrogen-limited media with 20 g L^-1^ glycerol as the sole carbon source for 72 h (Bhutada et al., 2017). Lipids were extracted from the cells (Bhutada et al., 2017) and the total FA and sn-2 FA composition of TAG were analysed (van Erp et al., 2019).

WT cells contained TAG with around 21% 16:0, 46% 18:1 and 14% 18:2 (Table 2). Palmitoleic acid (16:1) and stearic acid (18:0) and were also present at ∼6% and ∼12%, respectively (Table 2). As has been reported previously (Thorpe and Ratledge 1972), 16:0 (and 18:0) are virtually absent from the sn-2 position (Table 2). Only ∼3% of the 16:0 in TAG is present at the sn-2 position. Since we chose to integrate cassettes at the *GSY1* locus, we also analysed *gsy1Δ* cells as a control. The FA composition of TAG from *gsy1Δ* is altered, with a significant (P < 0.05) decrease in the percentage of 16:0 (Table 2), which has been reported previously (Bhutada et al., 2017). However, 16:0 was still excluded from the sn-2 position of TAG in *gsy1Δ* cells (Table 2). Cells of *gsy1Δ-mLPAT1* exhibited only small differences in the FA composition of TAG verses *gsy1Δ* and a small but significant (P < 0.05) increase in the percentage of 16:0 at sn-2 to ∼12%. By contrast, *gsy1Δ-AGPAT1* and *gsy1Δ-LPAAT2* cells contained TAG with both an increase in total 16:0 verses *gsy1Δ* and a large increase in 16:0 at the sn-2 position (P < 0.05) (Table 2). By comparing the total FA composition of TAG to that at sn-2, we calculated that in *gsy1Δ-AGPAT1* and *gsy1Δ-LPAAT2* cells respectively, ∼58% and ∼63% of the total 16:0 in TAG is at the sn-2 position. Of the strains we tested, *gsy1Δ-LPAAT2* has a FA composition that is closest to human milk fat (Lopez-Lopez et al., 2002) with ∼19% 16:0, ∼3% 16:1, ∼13% 18:0, ∼50% 18:1 and ∼15% 18:2, with more than 60% of the total 16:0 at the sn-2 position (Table 2).

**Table 2.** Total and sn-2 FA composition of TAG from *Y. lipolytica* strains cultured on 20 g L^-1^ glycerol in nitrogen-limited media. Values are means ±SD of measurements made on three separate cultures for each genotype. * denote values significantly (P < 0.05) different from WT (ANOVA + Tukey HSD test).

| Strain | 16:0 | 16:1 | 18:0 | 18:1 | 18:2 |
| --- | --- | --- | --- | --- | --- |
| *Total FA composition of TAG (%)* |  |  |  |  |  |
| WT | 21.3 ± 1.2 | 6.5 ± 0.5 | 11.7 ± 0.4 | 46.3 ± 2.4 | 14.2 ± 1.2 |
| *gsy1Δ* | 9.7 ± 0.9* | 4.7 ± 0.6* | 11.5 ± 0.8 | 58.0 ± 2.2* | 16.1 ± 1.4* |
| *gsy1Δ-mLPAT1* | 9.0 ± 0.2* | 3.7 ± 0.1* | 14.2 ± 0.1* | 60.5 ± 0.5* | 12.7 ± 0.2* |
| *gsy1Δ-AGPAT1* | 16.6 ± 0.6* | 3.6 ± 0.3* | 11.3 ± 0.3 | 52.8 ± 0.8* | 15.7 ± 0.5* |
| *gsy1Δ-LPAAT2* | 19.2 ± 1.3 | 3.3 ± 0.2* | 12.6 ± 4.1 | 50.3 ± 4.2 | 14.6 ± 1.8 |
| *FA composition (and percentage of total 16:0) at the sn-2 position of TAG (%)* |  |  |  |  |  |
| WT | 2.2 ± 1.3 (**3.5 ± 1.3**) | 2.1 ± 0.2 | 1.9 ± 1.9 | 73.1 ± 0.5 | 20.7 ± 2.4 |
| *gsy1Δ* | 1.6 ± 0.5 (**5.5 ± 2.1**) | 1.2 ± 0.5* | 0.6 ± 0.6 | 75.6 ± 1.5* | 21.0 ± 1.7 |
| *gsy1Δ-mLPAT1* | 3.1 ± 0.5 (**11.6 ± 1.0\***) | 1.1 ± 0.1* | 1.6 ± 0.4 | 77.2 ± 1.3* | 17.0 ± 0.3* |
| *gsy1Δ-AGPAT1* | 28.7 ± 0.5* (**57.8 ± 1.6\***) | 3.2 ± 0.1* | 1.4 ± 0.2 | 49.0 ± 0.4* | 17.7 ± 0.8 |
| *gsy1Δ-LPAAT2* | 35.9 ± 1.4* (**62.7 ± 3.6\***) | 3.3 ± 0.8* | 3.6 ± 3.8 | 43.7 ± 1.7* | 13.4 ± 1.2* |

### 3.2. Culture on palm oil or a mixture of glucose and palm oil improves TAG composition

*Y. lipolytica* can utilise a range of carbon sources in addition to glycerol, including various sugars and lipids (Spagnuolo et al., 2018). It is known that the carbon source can influence both the quantity of TAG that is produced and its FA composition in *Y. lipolytica* (Papanikolaou et al. 2003; Athenstaedt et al., 2006; Beopoulos et al., 2008; Vasiliadoua et al., 2018a, b). We therefore grew WT and *gsy1Δ-LPAAT2* in shake flask culture on nitrogen-limited media with 20 g L^-1^ glucose or 20 g L^-1^ palm oil as sole carbon sources, or with a mixture of 10 g L^-1^ of each substrate. We extracted lipids from cells at stationary phase (Bhutada et al., 2017) and determined the total FA and sn-2 composition of TAG (van Erp et al., 2019). The FA composition of TAG from WT and *gsy1Δ-LPAAT2* cells grown on glucose (Table 3) was similar with that of cells grown on glycerol (Table 2) except that 18:0 levels were significantly (P < 0.05) lower (∼8% verses ∼12%). The percentage of total 16:0 that is at the sn-2 position was only ∼1% in WT, whereas it was ∼62% in *gsy1Δ-LPAAT2* (Table 3). This is a similar level of 16:0 enrichment at sn-2 as we found for culture on glycerol (Table 2).

**Table 3.** Total and sn-2 FA composition of TAG from *Y. lipolytica* strains cultured on 20 g L^-1^ glucose in nitrogen-limited media. Values are means ±SD of measurements made on three separate cultures for each genotype. nd is not detected. * denote values significantly (P < 0.05) different from WT (Student’s t-tests).

| Strain | 16:0 | 16:1 | 18:0 | 18:1 | 18:2 |
| --- | --- | --- | --- | --- | --- |
| *Total FA composition of TAG (%)* |  |  |  |  |  |
| WT | 22.5 ±0.1 | 6.5 ±0.1 | 8.6 ±0.1 | 50.1 ±0.1 | 12.2 ±0.3 |
| *gsy1Δ-LPAAT2* | 19.8 ±0.3* | 3.5 ±0.1* | 8.3 ±0.1* | 58.3 ±0.1* | 10.1 ±0.2* |
| *FA composition (and percentage of total 16:0) at the sn-2 position of TAG (%)* |  |  |  |  |  |
| WT | 0.7 ±0.4 (**1.10 ±0.56**) | nd | 0.6 ±0.4 | 76.9 ±0.9 | 20.5 ±0.2 |
| *gsy1Δ-LPAAT2* | 36.5 ±0.3* (**61.6 ±0.94\***) | 0.8 ±0.1* | 0.4 ±0.1 | 49.0 ±0.5* | 10.4 ±0.2* |

The FA composition of TAG from WT and *gsy1Δ-LPAAT2* cells grown on palm oil (Table 4) was different from cells cultured on glycerol or glucose as sole carbon source (Table 2 and 3). The levels of 16:0 increased significantly (P < 0.05) to more than 26% and 16:1 and 18:0 reduced also to ∼4 and ∼1%, respectively (Fig 3a). These changes are consistent with the FA composition of the palm oil substrate, which is rich in 16:0, 18:1 and 18:2, but contains little 16:1 and 18:0 (Supplementary Table 2). The percentage of total 16:0 at the sn-2 position was only ∼1% in WT, but it increased to ∼62% in *gsy1Δ-LPAAT2* (Table 4), which is similar with the level of 16:0 enrichment at sn-2 obtained when this strain was cultured on glycerol or glucose (Table 2 and 3). However, total 16:0 content in TAG is higher, while 18:0 at the sn-1/3 positions is also decreased when *gsy1Δ-LPAAT2* cells are cultured on palm oil verses either glucose or glycerol, because the total level of 18:0 in TAG is lowered and 18:0 is predominantly esterified at the sn-1/3 positions (Table 4).

**Table 4.** Total and sn-2 FA composition of TAG from *Y. lipolytica* strains cultured on 20 g L^-1^ palm oil in nitrogen-limited media. Values are means ±SD of measurements made on three separate cultures for each genotype. nd is not detected. * denote values significantly (P < 0.05) different from WT (Student’s t-tests).

| Strain | 16:0 | 16:1 | 18:0 | 18:1 | 18:2 |
| --- | --- | --- | --- | --- | --- |
| *Total FA composition of TAG (%)* |  |  |  |  |  |
| WT | 30.3 ±0.5 | 5.4 ±1.1 | 2.2 ±0.2 | 39.2 ±0.3 | 22.8 ±0.4 |
| *gsy1Δ-LPAAT2* | 26.2 ±1.0* | 3.7 ±0.1* | 1.1 ±0.1* | 42.2 ±0.6* | 26.3 ±0.7* |
| *FA composition (and percentage of total 16:0) at the sn-2 position of TAG (%)* |  |  |  |  |  |
| WT | 0.40 ±0.1 (**0.5 ±0.1**) | nd | 0.3 ±0.1 | 64.0 ±0.3 | 35.3 ±0.2 |
| *gsy1Δ-LPAAT2* | 48.8 ±0.9* (**62.2 ±3.1\***) | 0.7 ±0.1* | 1.0 ±0.8 | 37.5 ±1.0* | 11.7 ±0.6* |

We also cultured WT and *gsy1Δ-LPAAT2* cells on a mixture of glucose and palm oil and found that the FA composition of the TAG was similar to culture on palm oil alone, with significantly (P < 0.05) elevated levels of 16:0 (∼24%) and lowered levels of 16:1 and 18:0 (∼3% and ∼1%) (Table 5) verses culture on glycerol or glucose (Table 2 and 3). The percentage of total 16:0 at the sn-2 position was ∼1% in WT and increased to ∼69% in *gsy1Δ-LPAAT2*. The enrichment of 16:0 at sn-2 when *gsy1Δ-LPAAT2* cells were cultured on a mixture of glucose and palm oil (Table 5) was therefore significantly (P < 0.05) higher than when the cells were cultured on glycerol, glucose, or palm oil individually (Table 2, 3 and 4). As with culture on palm oil (Table 4), the reduction in 18:0 content also provided lower 18:0 levels at sn-1/3, and there was a reduction in 16:1 (Table 5).

**Table 5.** Total and sn-2 FA composition of TAG from *Y. lipolytica* strains cultured on 10 g L^-1^ glucose + 10 g L^-1^ palm oil in nitrogen-limited media. Values are means ±SD of measurements made on three separate cultures for each genotype. nd is not detected. * denote values significantly (P < 0.05) different from WT (Student’s t-tests).

| Strain | 16:0 | 16:1 | 18:0 | 18:1 | 18:2 |
| --- | --- | --- | --- | --- | --- |
| *Total FA composition of TAG (%)* |  |  |  |  |  |
| WT | 24.5 ± 2.5 | 3.0 ± 0.3 | 2.3 ± 0.1 | 48.6 ± 1.9 | 21.7 ± 0.5 |
| *gsy1Δ-LPAAT2* | 23.9 ± 3.1 | 3.2 ± 0.1 | 1.2 ± 0.1* | 49.4 ± 2.7 | 21.7 ± 1.2 |
| *FA composition (and percentage of total 16:0) at the sn-2 position of TAG (%)* |  |  |  |  |  |
| WT | 0.6 ± 0.5 (**0.9 ± 0.7**) | nd | 0.5 ± 0.3 | 69.1 ± 1.0 | 29.8 ± 1.2 |
| *gsy1Δ-LPAAT2* | 49.4 ± 3.8* (**69.1 ± 4.0\***) | 0.7 ± 0.1 | 0.5 ± 0.1 | 39.0 ± 3.9* | 9.8 ± 0.1* |

### 3.3. Culture on palm oil or a mixture of glucose and palm oil increases lipid titre

To determine whether *LPAAT2* expression affects *Y. lipolytica* cell biomass and lipid production in shake flask culture in nitrogen limited media we grew WT and *gsy1Δ-LPAAT2* on glycerol, glucose, palm oil and a mixture of glucose and palm oil. After 72 hours of culture on these carbon sources the cells had reached stationary phase (Bhutada et al., 2017). On glycerol or glucose, WT cell biomass reached ∼4 g L^-1^ (Table 6), whereas a significantly (P < 0.05) higher biomass of ∼6 g L^-1^ was achieved on palm oil and on glucose plus palm oil (Table 6). *gsy1Δ-LPAAT2* biomass was significantly (P < 0.05) reduced when cells were cultured on glycerol and glucose but remained around the same as WT on palm oil or glucose plus palm oil (Table 6). The lipid content as a percentage of cell dry weight (% CDW) of *gsy1Δ-LPAAT2* cells grown on glycerol or glucose was significantly (P < 0.05) higher than WT (Table 6). This increase is likely attributable to disruption of the *GSY1* locus, rather than to *LPAAT2* expression. *GSY1* disruption is known to block glycogen synthesis and enhance lipid content (Bhutada et al., 2017). A comparison of *gsy1Δ* and *gsy1Δ-LPAAT2* cells grown on nitrogen limited media with glucose showed that *gsy1Δ* has a slightly higher lipid content (18.2 ±0.3 % verses 17.0 ±0.2 % (n=3)). The biomass of *gsy1Δ* and *gsy1Δ-LPAAT2* cells grown on glucose were 3.6 ±0.1 and 3.7 ±0.1 g L^-1^ (n=3) respectively, which are not significantly different values (P > 0.05) and this suggests that *LPAAT2* expression is not detrimental to growth on this carbon source. The lipid content of WT and *gsy1Δ-LPAAT2* cells grown on palm oil and on glucose plus palm oil was similar (in the range of 42 to 48%) (Table 6). These values are more than double those obtained from culture on glycerol or glucose as a sole carbon source (Table 6). The combined gains in cell biomass and lipid content associated with culture on palm oil or on glucose plus palm oil mean that the lipid titre (∼3 g L^-1^) is four to five time higher than for culture on glycerol or glucose (Table 6).

**Table 6.** Cell biomass and lipid content of *Y. lipolytica* strains cultured on 20 g L^-1^ glycerol, glucose, or palm oil or on 10 g L^-1^ glucose + 10 g L^-1^ palm oil in nitrogen-limited media. Values are means ±SD of measurements made on three separate cultures for each genotype. * denote values significantly (P < 0.05) different from WT (Student’s t-tests) and ^#^ from glycerol (ANOVA + Tukey HSD test).

| Strain | Substrate | Cell biomass<br>(g L<sup>-1</sup>) | Lipid content<br>(% of CDW) | Lipid titre<br>(g L<sup>-1</sup>) |
| --- | --- | --- | --- | --- |
| WT | Glycerol | 4.0 ±0.1 | 11.5 ±0.5 | 0.5 ±0.1 |
| *gsy1Δ-LPAAT2* |  | 3.3 ±0.2* | 21.5 ±0.9* | 0.7 ±0.1* |
| WT | Glucose | 4.0 ±0.1 | 10.1 ±1.8 | 0.4 ±0.1<sup>#</sup> |
| *gsy1Δ-LPAAT2* |  | 3.7 ±0.1*<sup>#</sup> | 17.0 ±1.7*<sup>#</sup> | 0.6 ±0.1*<sup>#</sup> |
| WT | Palm oil | 6.3 ±1.1<sup>#</sup> | 48.0 ±6.5<sup>#</sup> | 3.1 ±0.8<sup>#</sup> |
| *gsy1Δ-LPAAT2* |  | 6.7 ±1.3*<sup>#</sup> | 46.6 ±0.4<sup>#</sup> | 3.1 ±0.5<sup>#</sup> |
| WT | Glucose + Palm oil | 5.9 ±0.2<sup>#</sup> | 42.4 ±2.2<sup>#</sup> | 2.5 ±0.1<sup>#</sup> |
| *gsy1Δ-LPAAT2* |  | 5.9 ±1.3<sup>#</sup> | 46.8. ±2.1* | 2.8 ±0.5<sup>#</sup> |

### 3.4. LPAAT2 expression also leads to 16:0 enrichment at the sn-2 position in an obese strain

Several studies have showed that *Y. lipolytica* can be engineered to enhance TAG accumulation (Ledesma-Amaro and Nicaud, 2016). Lazar et al., (2014) created an ‘*obese*’ strain by overexpressing acyl-CoA:diacylglycerol acyltransferase (*DGA2*) and glycerol-3-phosphate dehydrogenase (*GPD1*) to enhance TAG biosynthesis (Beopoulos et al., 2012; Dulermo and Nicaud, 2011) and by deleting the six genes encoding acyl-coenzyme A oxidases (*POX1-6*) and the TAG lipase *TGL4* to block peroxisomal FA β-oxidation and TAG hydrolysis, respectively (Beopoulos et al., 2008; Dulermo et al., 2013). To determine whether *LPAAT2* expression also leads to 16:0 enrichment at the sn-2 position of TAG in this *obese* strain we integrated *TEF1p-LPAAT2* at the *GSY1* locus to create the strain *obese-gsy1Δ-LPAAT2* (Table 1). Both the *obese* and *obese-gsy1Δ-LPAAT2* strains were grown in triplicate in shake flask cultures in nitrogen-limited media with 20 g L^-1^ glucose or with 10 g L^-1^ glucose plus 10 g L^-1^ palm oil as the carbon source. Palm oil was not tested as a sole carbon source because the *obese* strain is deficient in FA β-oxidation (Beopoulos et al., 2008).

The total FA composition of TAG from *obese* cells grown on glucose as a sole carbon source differs from that of WT, in that 16:0 is increase to ∼30% and 18:2 is reduced to ∼6% (Lazar et al., 2014). The percentage of 16:0 in TAG from *obese-gsy1Δ-LPAAT2* cells is significantly (P < 0.05) lower than in *obese* on both glucose (Table 7) and on glucose plus palm oil (Table 8). This is likely caused by disruption of *GSY1* (Bhutada et al., 2017). Growth of *obese* on glucose plus palm oil also leads to a significantly (P < 0.05) lower percentage of 16:1 and 18:0 (Table 8), as was observed in WT (Table 5). Less than ∼1% of the 16:0 in TAG is present at the sn-2 position in the *obese* strain grown either on glucose or on glucose plus palm oil (Table 7 and 8). In TAG from *obese-gsy1Δ-LPAAT2* cells, ∼52% and ∼62% of the 16:0 is at the sn-2 position when the cells are grown on glucose and on glucose plus palm oil, respectively (Table 7 and 8). Enrichment of 16:0 at the sn-2 position therefore appears to be a bit lower in *obese-gsy1Δ-LPAAT2* than we measured in *gsy1Δ-LPAAT2*, but culture on glucose plus palm oil also led to a significant (P < 0.05) increase verses glucose alone (Table 7 and 8).

**Table 7.** Total and sn-2 FA composition of TAG from *Y. lipolytica obese* strains cultured on 20 g L^-1^ glucose in nitrogen-limited media. Values are means ±SD of measurements made on three separate cultures for each genotype. * denote values significantly (P < 0.05) different from *obese* (Student’s t-tests).

| Strain | 16:0 | 16:1 | 18:0 | 18:1 | 18:2 |
| --- | --- | --- | --- | --- | --- |
| *Total FA composition of TAG (%)* |  |  |  |  |  |
| *obese* | 30.4 ±0.4 | 6.6 ±0.2 | 10.3 ±0.1 | 46.5 ±0.5 | 6.1 ±0.1 |
| *obese-gsylA-*<br>*LPAAT2* | 26.2 ±0.3* | 4.2 ±0.1* | 9.5 ±0.2* | 52.9 ±0.6* | 7.2 ±0.3* |
| *FA composition (and percentage of total 16:0) at the sn-2 position of TAG (%)* |  |  |  |  |  |
| *obese* | 0.2 ±0.1 (**0.2 ±0.1**) | 2.6 ±0.2 | 0.2 ±0.1 | 85.2 ±1.4 | 11.7 ±0.7 |
| *obese-gsylA-*<br>*LPAAT2* | 40.6 ±0.7* (**51.6 ±0.5\***) | 6.3 ±0.1* | 1.0 ±0.2* | 44.3 ±0.8* | 7.8 ±0.6* |

**Table 8.** Total and sn-2 FA composition of TAG from *Y. lipolytica obese* strains cultured on 10 g L^-1^ glucose + 10 g L^-1^ palm oil in nitrogen-limited media. Values are means ±SD of measurements made on three separate cultures for each genotype. nd is not detected. * denote values significantly (P < 0.05) different from *obese* (Student’s t-tests).

| Strain | 16:0 | 16:1 | 18:0 | 18:1 | 18:2 |
| --- | --- | --- | --- | --- | --- |
| *Total FA composition of TAG (%)* |  |  |  |  |  |
| *obese* | 31.5 ±0.5 | 2.8 ±0.2 | 2.2 ±0.1 | 51.8 ±1.9 | 11.7 ±0.5 |
| *obese-gsylA-*<br>*LPAAT2* | 27.1 ±0.4* | 3.1 ±0.3 | 1.4 ±0.1* | 54.7 ±2.7* | 13.7 ±1.2* |
| *FA composition (and percentage of total 16:0) at the sn-2 position of TAG (%)* |  |  |  |  |  |
| *obese* | 0.8 ±0.5 (**1.0 ±0.6**) | nd | 0.6 ±0.1 | 77.8 ±1.0 | 20.8 ±1.2 |
| *obese-gsylA-*<br>*LPAAT2* | 50.4 ±2.8* (**62.0 ±3.1\***) | 0.9 ±0.1 | 0.7 ±0.1 | 38.0 ±2.5* | 10.0 ±0.3* |

Using glucose as sole carbon source, *obese* and *obese-gsy1Δ-LPAAT2* cell biomass reached around 5.8 g L^-1^, whereas a significantly (P < 0.05) higher biomass of ∼7.5 g L^-1^ was achieved on glucose plus palm oil (Table 9). The lipid content, as a % CDW, of *obese* and *obese-gsy1Δ-LPAAT2* cells grown on glucose was ∼41% and ∼46%, respectively (Table 9). The increase in *obese-gsy1Δ-LPAAT2* is likely attributable to the disruption of *GSY1* (Bhutada et al., 2017). On glucose plus palm oil, *obese* and *obese-gsy1Δ-LPAAT2* lipid content reached around ∼67% of CDW. The gains in cell biomass and lipid content associated with culture on glucose plus palm oil mean that the lipid titre (∼5.3 g L^-1^) is nearly two times higher than for culture on glucose (Table 9). Lipid titre in *obese-gsy1Δ-LPAAT2* was also substantially higher than in *gsy1Δ-LPAAT2*, which is consistent with the fact that the *obese* strain was previously engineered to accumulate more lipid (Lazar et al., 2014).

**Table 9.** Cell biomass and lipid content of *Y. lipolytica obese* strains cultured on 20 g L^-1^ glucose, or on 10 g L^-1^ glucose + 10 g L^-1^ palm oil in nitrogen-limited media. Values are means ±SD of measurements made on three separate cultures for each genotype. * denote values significantly (P < 0.05) different from *obese* and ^#^ from glucose (Student’s t-tests).

| Strain | Substrate | Cell biomass<br>(g L<sup>-1</sup>) | Lipid content<br>(% of CDW) | Lipid titre<br>(g L<sup>-1</sup>) |
| --- | --- | --- | --- | --- |
| *obese* | Glucose | 5.8 ±0.3 | 41.5 ±0.8 | 2.4 ±0.2 |
| *obese-gsy1Δ-LPAAT2* |  | 5.5 ±0.2* | 45.9 ±1.7* | 2.5 ±0.4 |
| *obese* | Glucose + Palm oil | 7.5 ±0.2<sup>#</sup> | 67.2 ±3.2<sup>#</sup> | 5.0 ±0.2<sup>#</sup> |
| *obese-gsy1Δ-LPAAT2* |  | 7.7 ±0.5<sup>#</sup> | 68.9 ±2.0<sup>#</sup> | 5.3 ±0.3<sup>#</sup> |

## 4. Discussion

In this study we show that *Y. lipolytica* cells, that normally exclude 16:0 from the sn-2 position in their TAG (Thorpe and Ratledge 1972), can be engineered to incorporate more than 60% of the total 16:0 at the sn-2 position by expressing a LPAT with a preference of 16:0-CoA. The most effective LPAT we tested is LPAAT2 from the alga *Chlamydomonas reinhardtii* (Kim et al., 2018). The change in TAG stereoisomeric structure, combined with the native FA composition found in *Y. lipolytica* cells cultured on glycerol or glucose, produces fat that is similar to human milk, in terms of the major molecular species of FAs (16:0, 18:1 and 18:2), and thus has the potential to be used as a HMFS (β-palmitate) ingredient in infant formulas. Furthermore, we found that both the FA composition and yield of lipid that is produced by wildtype and a lipid overproducing *obese Y. lipolytica* strain (Lazar et al., 2014) expressing *LPAAT2* can be improved, without further metabolic engineering, by culture on palm oil or on a mixture of glucose plus palm oil. Unexpectedly the combination of glucose plus palm oil also led to an increase in the percentage of 16:0 in the sn-2 position up to ∼70%, which is similar to values reported for human milk fat (Breckenridge et al., 1969; Giuffrida et al., 2018). The reason why this mixture of substrates leads to a greater enrichment of 16:0 at sn-2 is unclear but combinations can influence the regulation of metabolism (Fabiszewska et al., 2015; Lubuta et al., 2019). *Y. lipolytica* uses glucose initially as a preferred carbon source when cultured on glucose plus the free FA 18:1, and then switches to use 18:1 once the glucose is depleted (Kamzolova et al., 2011).

*Y. lipolytica* is known to be flexible with regards to its carbon source and (as its name suggests) it can grow readily on a range of lipids as well as on glycerol, acetate, and certain sugars (Fickers et al., 2005: Spagnuolo et al., 2018). Its substrate range has also been successfully extended by GE (Spagnuolo et al., 2018), for example to include lignocellulosic materials (Niehus et al., 2018). The highest productivity in terms of lipids has been reported for GE strains in nitrogen-limited glucose-fed batch culture, where 1.2 g^-1^ L^-1^ h^-1^ has been achieved (Qiao et al., 2017). It is noteworthy that similar or higher lipid productivities have also been reported for *Rhodosporidium toruloides, Rhodotorula glutinis* and *Lipomyces starkeyi* yeasts cultivated on glucose (Karamerou et al., 2021). The *Y. lipolytica* strain used in this study has not been engineered to improve its total FA composition, but modification of this trait is possible based on the findings of previous studies (Ledesma-Amaro and Nicaud, 2016; Tsakraklides et al., 2018). It is noteworthy that human milk fat also contains low levels of medium chain saturated FAs (∼12%) and very long chain polyunsaturated FAs (∼2%) that are both absent from *Y. lipolytica* (Lopez-Lopez et al., 2002; Carsanba et al., 2020). These FAs are incorporated into the fat phase of infant formulas from palm kernel or coconut oil and from algae or fish oils, respectively (Wei et al., 2019). However, both these classes of FA can also be produced in *Y. lipolytica* by metabolic engineering (Rutter et al., 2015; Xue et al., 2013). Further enrichment of 16:0 at the sn-2 position in TAG may also be achievable in *Y. lipolytica* using GE, based on our experience in engineering this trait in oilseeds (van Erp et al., 2019; 2021).

*Y. lipolytica* produces extracellular lipases that hydrolyse lipids, allowing the cells to take up the free FAs and β-oxidise them (Fickers et al., 2005). The FAs are also used for lipid synthesis and, when in excess, are incorporated into TAG (Fickers et al., 2005). The FA composition of the TAG therefore partially reflects that of the lipid substrate (Papanikolaou et al. 2003; Vasiliadoua et al., 2018a, b). The palm oil that we used as a substrate in this study has a FA composition that differs from *Y. lipolytica*, leading to the synthesis of TAG that is closer to a HMFS. Specifically, the palm oil contributes a higher 16:0 content and a lower 18:0 and 16:1 content in *Y. lipolytica* TAG. *Y. lipolytica* also discriminates against uptake of exogenous 18:0 but selectively incorporates it into TAG (Papanikolaou and Aggelis 2003). Human milk fat contains both 16:1 and 18:0 (Lopez-Lopez et al., 2002) but they are less abundant than in wildtype (W29) *Y. lipolytica* cultured on glycerol or glucose. 18:0 is also mainly esterified to the sn-1/3 positions in *Y. lipolytica* TAG; even in cells expressing *LPAAT2* or *AGPAT1*. These LPATs either exhibit fatty acyl chain length specificity that prohibits them from incorporating 18:0 into the sn-2 position of glycerolipids or they have comparatively little 18:0-CoA substrate. It is noteworthy that most of the 18:0 that is present in human milk fat is in the sn-1/3 positions, unlike 16:0 (Lopez-Lopez et al., 2002). However, it is likely desirable to excluded both 16:0 and 18:0 from the sn-1/3 positions of TAG used as a HMFS, because when either of these long chain saturated FAs are released by sn-1/3 specific lipases in the infant gut they can form calcium soaps and aggregate, reducing lipid and calcium absorption (Innis 2011; Béghin et al., 2018).

It is known that when wildtype and lipid overproducing *Y. lipolytica* strains are cultured on lipids or on lipid and sugar mixtures the cells can achieve a higher lipid content (% CDW) than when the cells are cultured on sugars alone (Beopoulos et al., 2008). Papanikolaou et al., (2003) investigated the capacity of a wildtype *Y. lipolytica* strain to produce cocoa butter equivalent (another commercially important class of structured lipid) using animal stearin, and reported that culture on stearin plus glycerol yielded the highest titre of ∼3.4 g L^-1^, which is similar to the titre we obtained through shake flask culture of *gsy1Δ-LPAAT2* on palm oil or on glucose plus palm oil. The lipid content we measured (42 to 48% of CDW), is also similar with that reported by Papanikolaou et al., (2003). However, it is higher than the values reported by Vasiliadoua et al., (2018a, b) who also used palm oil as a sole carbon source. In the *obese* genetic background cultured on glucose as sole carbon source, lipid titre and lipid content (% CDW) are increased (Lazar et al., 2014). We found that this is also the case for culture on glucose plus palm oil. It is noteworthy that all the genetic modifications introduced into the *obese* background are designed to increase TAG accumulation, rather than directly targeting FA synthesis (Lazar et al., 2014). Integration of *TEFp-LPAAT2* at the *GSY1* locus also disrupts glycogen synthesis, which results in higher TAG content (Bhutada et al., 2017). The *obese-gsy1Δ-LPAAT2* strain is therefore appropriately adapted to enhance lipid production using a lipid substrate.

Some mammals (e.g. *H. sapiens*), algae (e.g. *Nannochloropsis oceanica* and *C. reinhardtii*) and bacteria (e.g. *Rhodococcus opacus*) can incorporate 16:0 into the sn-2 position of their TAG (Breckenridge et al., 1969; Nobusawa et al., 2017; Kim et al., 2018; Wältermann et al., 2000). The algae and bacteria that are known to do this have FA compositions that are rather dissimilar to human milk fat. *R. opacus* for instance produces a high percentage of odd chain length FAs when cultured on glucose or glycerol (Kurosawa et al., 2010; 2015). However, *R. opacus* is lipolytic and the FA composition of its TAG can also reflect that of its lipid substrate (Alvarez et al., 1996). Zang et al., (2020a, b, c) have recently showed that HMFS can be made by culturing a wildtype *R. opacus* strain on mixtures of vegetable, animal and algal oils that have been chemically inter-esterified or on mixtures of FA ethyl esters. However, the lipid titre they report in shake flask culture under nitrogen limitation (Zang et al., 2020a, b, c) are around a half and a quarter of the values we report here using an engineered *Y. lipolytica* strain grown on glucose or on glucose plus palm oil, respectively.

There are several issues to consider when using lipid (as opposed to sugar) substrates for microbial TAG production. For example, substrates that are immiscible in water such as lipids require emulsification and may therefore be harder to work with in large-scale aerobic fermenters. More of the neutral lipid that accumulates in the cells can also be in the form of sterol esters and free FAs (Athenstaedt et al., 2006; Beopoulos et al., 2008). Vegetable oils (and their chemically modified derivatives) are also more expensive substrates than sugars (weight for weight), but they do have a higher energy density and the carbon conversion efficiency of converting lipids to lipids is greater, theoretically. Ultimately, a thorough technoeconomic analysis is required to determine what the lowest cost microbial fermentation process would likely be for HMFS production. However, for heterotrophic growth of oleaginous yeast on sugar substrates, many analyses have already been made that may be applicable, owing to interest in microbial production of similar lipids such as palm oil substitute (Karamerou et al., 2021). Karamerou et al., (2021) recently estimated that the lowest selling cost it may be theoretically possible to reach is ∼$1.2 kg^-1^ for a production scale of 48k tonnes yr^-1^. Market research suggests that HMFS production is on a similar scale and the price exceeds ∼$6 kg^-1^. Finally, on environmental and sustainability grounds, it may ultimately be considered preferable to use sugars, rather than vegetable oils (and palm oil in particular), as substrates for HMFS production.

## Authorship contributions

P.J.E. conceived the research and supervised the experiments; G.B. designed and performed the experiments and analysed the data, with some assistance from G.M. and P.J.E. R.L-A. provided materials and technical advice on cloning and transformation. P.J.E. wrote the article with contributions of all the authors.

## Acknowledgments

We wish to thank Dr Harrie van Erp for providing training in lipid analysis methods and Prof. Klaus Natter for providing materials.

