## Supplemental data for "Production of human milk fat substitute by engineered strains of *Yarrowia lipolytica*"

**Supplementary data.**

**Supplementary Figure 1.** Codon optimised LPAT sequences use in this study.

```
>mLPAT1
AAGCTTATGTCCGACCTGTCTGGTGCTGCTACCCCCGAGTCCACCTACCCTGAGCCTGAGATCAAGCTGTCCTCTCGA
CTCCGAGGAATTTGCTTCTGTCTCGTCGCCGGCGTGTCTGCTATCGTCCTGATTGTGCTCATGATCACCGGCCACCCC
TTCGTCCTGCTCTTCGACCGATACCGACGAAAGTTCCACCACTTCATCGCCAAGCTGTGGGCTTCCATCTCTATCTAC
CCCTTCTACAAGACCGACATTCAGGGTCTGGAGAACCTCCCCTCCTCTGACACCCCCTGCGTCTACGTGTCCAACCAC
CAGTCTTTCCTGGACATCTACACCCTGCTCTCCCTCGGACAGTCTTACAAGTTCATTTCCAAGACCGGCATCTTCGTC
ATTCCCGTGATCGGCTGGGCCATGTCCATGATGGGTGTCGTGCCCCTGAAGCGAATGGACCCCCGATCTCAGGTCGAC
TGCCTGAAGCGATGTATGGAGCTCGTCAAGAAGGGTGCCTCCGTCTTCTTCTTCCCCGAGGGAACCCGATCTAAGGAC
GGACGACTGGGCCCCTTCAAGAAGGGCGCTTTCACCATTGCTGCTAAGACCGGTGTGCCTGTGGTGCCCATTACCCTG
ATGGGCACCGGCAAGATCATGCCCACCGGTTCCGAGGGAATTCTCAACCACGGTGACGTCCGAGTGATCATTCACAAG
CCCATCTACGGATCTAAGGCTGACCTGCTCTGTGACGAGGCCCGAAACAAGATTGCTGAGTCCATGAACCTGCTCTCT
TAACGATCGTTTTTTTTTATATATATATATATATATATATATAACTGTCTAGAAATAAAGAGTATCATCTTTCAAAAAGC
TT

>AGPAT1
AAGCTTATGGACCTGTGGCCCGGAGCTTGGATGCTGCTCCTGCTCCTGTTCCTCCTCCTGCTGTTCCTCCTGCCCACC
CTGTGGTTCTGCTCCCCCTCTGCTAAGTACTTCTTCAAGATGGCCTTCTACAACGGTTGGATTCTGTTCCTGGCCGTC
CTGGCTATTCCCGTCTGTGCTGTGCGAGGACGAAACGTGGAGAACATGAAGATCCTCCGACTGATGCTCCTGCACATC
AAGTACCTGTACGGAATTCGAGTTGAGGTCCGAGGCGCCCACCACTTCCCTCCCTCCCAGCCTTACGTCGTGGTCTCT
AACCACCAGTCCTCTCTGGACCTCCTGGGTATGATGGAGGTGCTCCCTGGACGATGTGTCCCTATCGCTAAGCGAGAG
CTGCTCTGGGCTGGTTCCGCTGGACTGGCTTGTTGGCTGGCTGGCGTCATCTTCATTGACCGAAAGCGAACCGGTGAC
GCTATTTCCGTGATGTCTGAGGTGGCTCAGACCCTCCTGACCCAGGACGTTCGAGTCTGGGTGTTCCCTGAGGGAACC
CGAAACCACAACGGTTCCATGCTGCCCTTCAAGCGAGGCGCCTTCCACCTCGCTGTCCAGGCTCAGGTCCCTATTGTG
CCCATTGTCATGTCCTCTTACCAGGACTTCTACTGCAAGAAGGAGCGACGATTCACCTCTGGACAGTGTCAGGTCCGA
GTGCTGCCTCCCGTGCCTACCGAGGGACTGACCCCCGACGACGTTCCTGCTCTGGCTGACCGAGTCCGACACTCCATG
CTGACCGTGTTCCGAGAGATTTCTACCGACGGTCGAGGCGGTGGAGACTACCTCAAGAAGCCCGGCGGTGGAGGCTAA
CGATCGTTTTTTTTTATATATATATATATATATATATATAACTGTCTAGAAATAAAGAGTATCATCTTTCAAAAAGCTT

>LPAAT2
AAGCTTATGTCTGTCCTCACCAAGTGGCTGGGTCTCCCCTCTTTCCTGTTCTCCGTCTTCGTGTTCTACTGGTCTCTC
CCCATCTTCGCCATTCTGTACCGAATCCGATTCGCTTCCCTGGGAAAGCGAAACGACATGCTCGACTGGGCTCGAGCC
CTGGTCGCCTACTTCCGAGTGACCCTGCTCCAGGCTGGCGAGCACACCCTGTACAAGGGCGGTCCCTGCCTGTACCTC
TGTAACCACCGATCCTGGGCTGACTTCTTCATTGACGCTTACCTGACCGAGGGACGAGCTGCTCTCATGTCTCGATGG
CTGGTCTACTTCGTGTTCCCCGTCTTCTGCACCTCCTGTATGATCCTCAAGGGTATTGTCCTGTTCAAGCGAGGAACC
ATTGCCGACAAGGAAGCCTTCAACGCCTGGCTGGACCAGACCCTGGGATCCTCTCACGTCCCTGGACTGCTGGTGTAC
CCCGAGGGACACCGATCTACCAAGCCTGCCTCCCTGCCTCTCAAGCGAGGTATGCTCCACTACGCTCACTCTCGAAAG
CTGCCCGTGCAGATTGTCGTGACCCGAGGCAAGGACGAGGTCCTGTCCGAGAAGTCTCAGTCCGTGCACTTCGGACGA
ACCTGCGTCACCACCTTCTCTAAGGTGCTCAAGTCCGCTGACTACCCCAACTTCGAGGCCTTCTTCACCGACCTGCAG
GCTACCTGGGACTCTTGTTGGGCCGCTACCTACGGACTGGAGGACCTCAAGAACGTGCCTCGATTCTCTATGCCCGGA
CCTCAGGCCTACTCCTACTCCTCTTCCATGTGGGTGCAGCAGCTCGCCATCACCCTCGTGTCTATTCTGGTCTTCGCT
GGAGTTTGTTACGGCTCCTGGCGAGGTCTGGCCGCTGCCCTGGCTGCTACCGGTGCTGCCCAGCAGGTGGTTGCTCTG
GTGCTGGCTGCTTGGGTGGGTTCTTCCGTGCTCCGATCCTTCCTGTAACGATCGTTTTTTTTTATATATATATATATA
TATATATAACTGTCTAGAAATAAAGAGTATCATCTTTCAAAAAGCTT
```

**Supplementary Table 1.** Plasmids and PCR primers used in this study.

| Plasmid | Genotype | Source/description |
|---|---|---|
| *pFA6aURA3-09* | *Parent Plasmid (5'_MCS-loxP-PPURA3TT-loxP-MCS_3')* | Bhutada et al., (2017) |
| *pGMKGSY12* | *YlGSY1*$^{P}$*-loxP-PURA3T-loxP-YlGSY1*$^{T}$ | Bhutada et al., (2017) |
| *YlmLPAT1* | *pUC57-5'_mLPAT1-Syn*$^{T}$*_3'* | This work |
| *YlAGPAT1* | *pUC57-5'_AGPAT1-Syn*$^{T}$*_3'* | This work |
| *YlLPAAT2* | *pUC57-5'_LPAAT2-Syn*$^{T}$*_3'* | This work |
| *pGSYTEF* | *YlGSY1*$^{P}$*-TEF1*$^{P}$*-loxP-PURA3T-loxP-YlGSY1*$^{T}$ | This work |
| *pTEF-mLPAT1* | *YlGSY1*$^{P}$*-TEF1*$^{P}$*-mLPAT1-Syn*$^{T}$*-loxP-PURA3T-loxP-YlGSY1*$^{T}$ | This work |
| *pTEF-AGPAT1* | *YlGSY1*$^{P}$*-TEF1*$^{P}$*-AGPAT1-Syn*$^{T}$*-loxP-PURA3T-loxP-YlGSY1*$^{T}$ | This work |
| *pTEF-LPAAT2* | *YlGSY1*$^{P}$*-TEF1*$^{P}$*-LPAAT2-Syn*$^{T}$*-loxP-PURA3T-loxP-YlGSY1*$^{T}$ | This work |

| Primers | Sequence |
|---|---|
| *TEF-GSY-F* | CTCGCAACAACCGATTCCAACAAGAGACCG GGTTGGCGGCGCA |
| *TEF-GSY-R* | ATAACTTCGTATAATGTATGCTATACGAAGT TATAAGCTTTGAATGATTCTTATACTCAGAA GGAAATGCTTAA |
| *GSY1*$^{P}$*-F* | GAGGAGCTGTTGGAGGTACGC |

| | |
|---|---|
| *GSY1*$^{T}$*-R* | GAACATGTGTGCGTTTTCACTTTCG |
| *mLPAT1_R* | GTTCTCCAGACCCTGAATGTCG |
| *AGPAT1-R* | GACCTCAACTCGAATTCCGTAC |
| *LPAAT2-R* | GCCATCGAGACATGAGAGCAGC |

**Supplementary Table 2.** Total FA composition of palm oil use as a substrate in this study.

| FA | 14:0 | 16:0 | 18:0 | 16:1 | 18:1 | 18:2 | 18:3 |
|---|---|---|---|---|---|---|---|
| % | 0.7 ±<0.1 | 37.3 ±0.5 | 4.2 ±0.2 | nd | 35.5 ±1.0 | 10.4 ±0.4 | 0.6 ±<0.1 |

Values are means ±SD of measurements made on three samples. nd is not detected.
